# Improved flux profiling in genome-scale modeling of human cell metabolism

**DOI:** 10.1101/2025.05.13.653488

**Authors:** Cyriel Huijer, Xiang Jiao, Yun Chen, Rosemary Yu

## Abstract

Understanding human cell metabolism through genome-scale flux profiling is of interest to diverse research areas of human health and disease. Metabolic modeling using genome-scale metabolic models (GEMs) has the potential to achieve this, but has been limited by a lack of appropriate input data as model constraints. Here we show that GEM-based flux profiling simulations can be improved with an appropriate input data collection procedure and exo-metabolite exchange flux calculation method, called regression during exponential growth phase (REGP). Our results show that the GEM-simulated feasible flux space is constrained to a biologically meaningful region, allowing an exploration of the basic organizing principles of the feasible flux space. These improvements help to fulfil the promise of GEMs as a valuable tool in the study of human metabolism and future development of translational applications.

## Introduction

Metabolism of human cells is a highly complex network of thousands of metabolites and reactions. Alterations in cell metabolism are associated with many complex health conditions such as diabetes, inflammatory diseases, and cancer (1,2). Importantly, the defining feature of metabolism is not the concentrations of metabolites in the cell, but the metabolic fluxes (*r*) through reactions and pathways (3,4). Intracellular metabolic fluxes can be experimentally determined through isotope-labeled substrate tracing for a small subset of reactions (5,6), but to systematically profile all fluxes of a cell at the genome-scale, mathematical modeling is necessary.

Genome-scale metabolic models (GEMs) is a modeling framework wherein the complete metabolic network of a cell is reconstructed *in silico* (5,7). GEMs can be used for simulations to calculate the optimal (max or min) fluxes of each reaction, and determine the feasible flux space for the entire metabolic network, using techniques called flux balance analysis (FBA) and flux variability analysis (FVA) (8). FBA and FVA requires a small amount of input data as constraints, typically consisting of measured exchange fluxes (± measurement error) of a small number of exo-metabolites, such as glucose, lactate, and amino acids. With these experimentally measured input data, GEM simulations have been shown to be strikingly accurate in microorganisms such as *E. coli* and *S. cerevisiae* (9–11). Critically, FBA and FVA assume that cells are in steady-state. Thus, input data for these successful applications of GEM simulations have all been collected during exponential growth.

Building on the success of GEM simulations in microbial applications, there is considerable interest in studying human cell metabolism using Human-GEM (3,12,13). The current practice to determine exo-metabolite exchange fluxes in human cells is to use the consumption and release (CORE) (4) method (Fig 1A). In this method, exo-metabolite concentrations in the cell culture media are measured at a single time point, and exchange fluxes are then calculated based on the difference between the measured ‘spent’ media and the unused (‘fresh’) media (Fig 1A). Thus, CORE-calculated exchange fluxes are not true steady-state exchange fluxes, and the use of these values as constraints for FBA and FVA should be done with caution.

**Figure 1.**
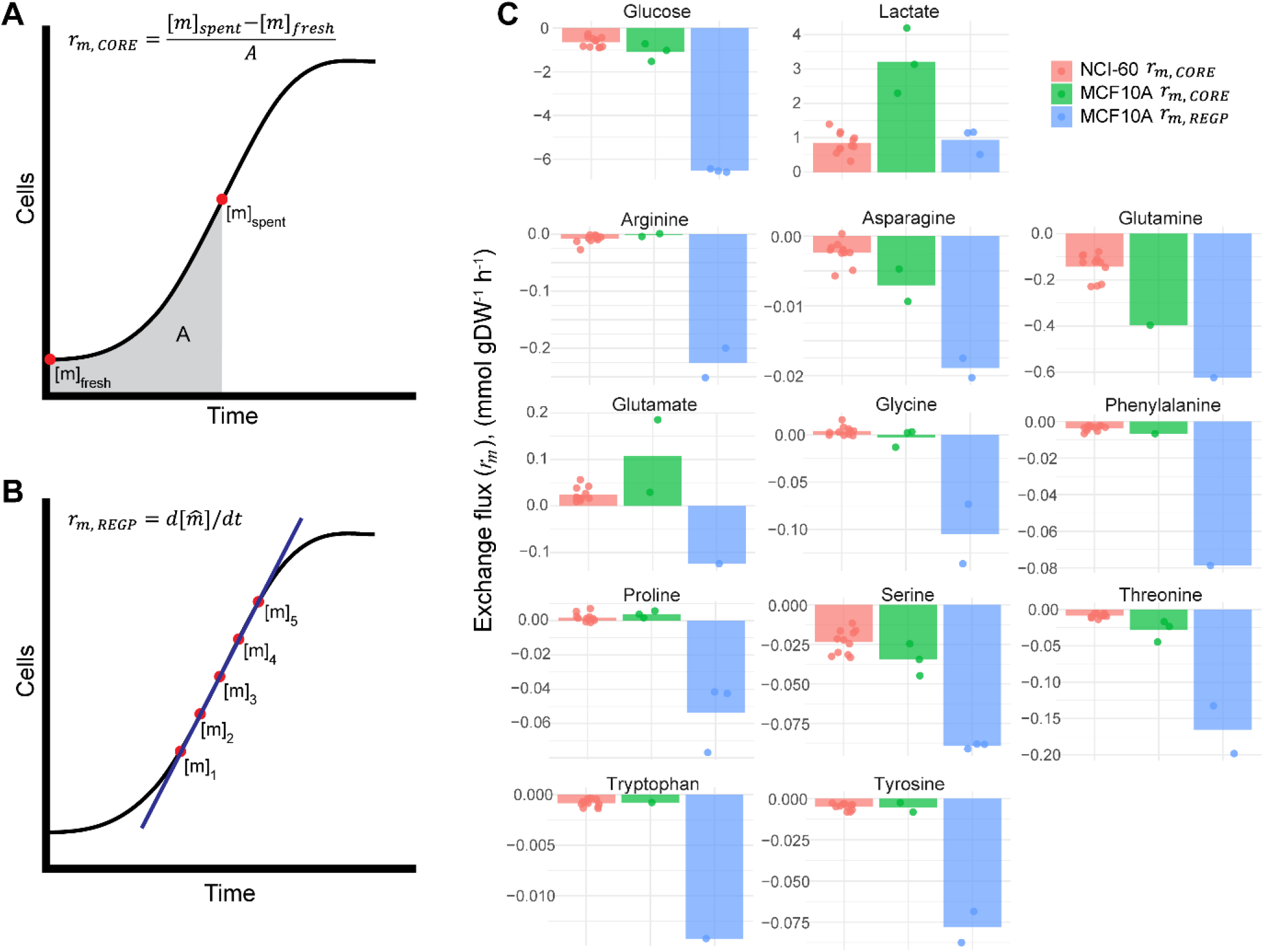
Exo-metabolite exchange fluxes. (A-B), schematic overview of exo-metabolite exchange flux calculation by the CORE and REGP methods. [*m*], exo-metabolite concentration; A, area under the curve; 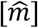, linear-regression-fitted exo-metabolite concentration (see Methods section). (C), exo-metabolite exchange fluxes (*r*_*m*_) for 11 cell lines in the NCI-60 panel calculated by the CORE method, and for the MCF10A cell line calculated by either the CORE or the REGP method.

In this study, we measured exo-metabolite concentrations at multiple time points during exponential growth for a human cell line MCF10A, and calculated exo-metabolite exchange fluxes by regression during exponential growth phase (REGP; Fig 1B). We found that REGP-calculated exchange fluxes (*r*_*m*,*REGP*_) were substantially different from CORE-calculated exchange fluxes (*r*_*m*,*CORE*_). Using *r*_*m*,*REGP*_ as input data for FBA and FVA, we showed that the GEM-simulated feasible flux space was constrained to a more biologically meaningful region, allowing an exploration of the basic organizing principles of the feasible flux space. We anticipate that future application of *r*_*m*,*REGP*_ as input data for GEM simulations can rapidly advance our understanding of cell metabolism in diverse applications related to human health and disease.

## Results

### Exo-metabolite exchange fluxes at steady-state

We measured the exo-metabolite concentrations of exponentially-growing MCF10A cells at five time points during the exponential growth steady-state (Supplemental Table 1), and used both the CORE (Fig 1A) and REGP (Fig 1B) methods to calculate the exchange fluxes of exo-metabolites, *r*_*m*_. Fig 1C shows the comparison between *r*_*m*,*REGP*_ and *r*_*m*,*CORE*_ in the MCF10A cell lines, as well as the *r*_*m*,*CORE*_ of 11 cell lines of the NCI-60 panel that were previously considered reliable (3,4,14). By convention, consumption of metabolites (e.g. glucose) is represented by a negative flux, and release of metabolites (e.g. lactate) by a positive flux. As expected, *r*_*m*,*CORE*_ were comparable between MCF10A cells and the 11 cell lines of the NCI-60 panel (Fig 1C). However, the *r*_*m*,*REGP*_ and *r*_*m*,*CORE*_ in MCF10A cells, based on the same raw metabolite measurements and cell count data, were substantially different. For several exo-metabolites, for example glutamate and glycine, *r*_*m*,*CORE*_ values were positive, indicating that cells were releasing these metabolites into the culture media; while *r*_*m*,*REGP*_ values were negative, indicating that cells were consuming these metabolites as nutrients. As the CORE method encompasses both lag phase and exponential growth phase (Fig 1A), whereas the REGP method calculates the exchange flux during exponential growth only (Fig 1B), this reflects that cell metabolism differs between lag phase and exponential growth (Fig 2). Our results indicate that glutamate is released by the cells during lag phase, and consumed during exponential growth (Fig 2B). Similarly, consumption of glycine differs between lag phase and exponential growth (Fig 2C).

**Figure 2.**
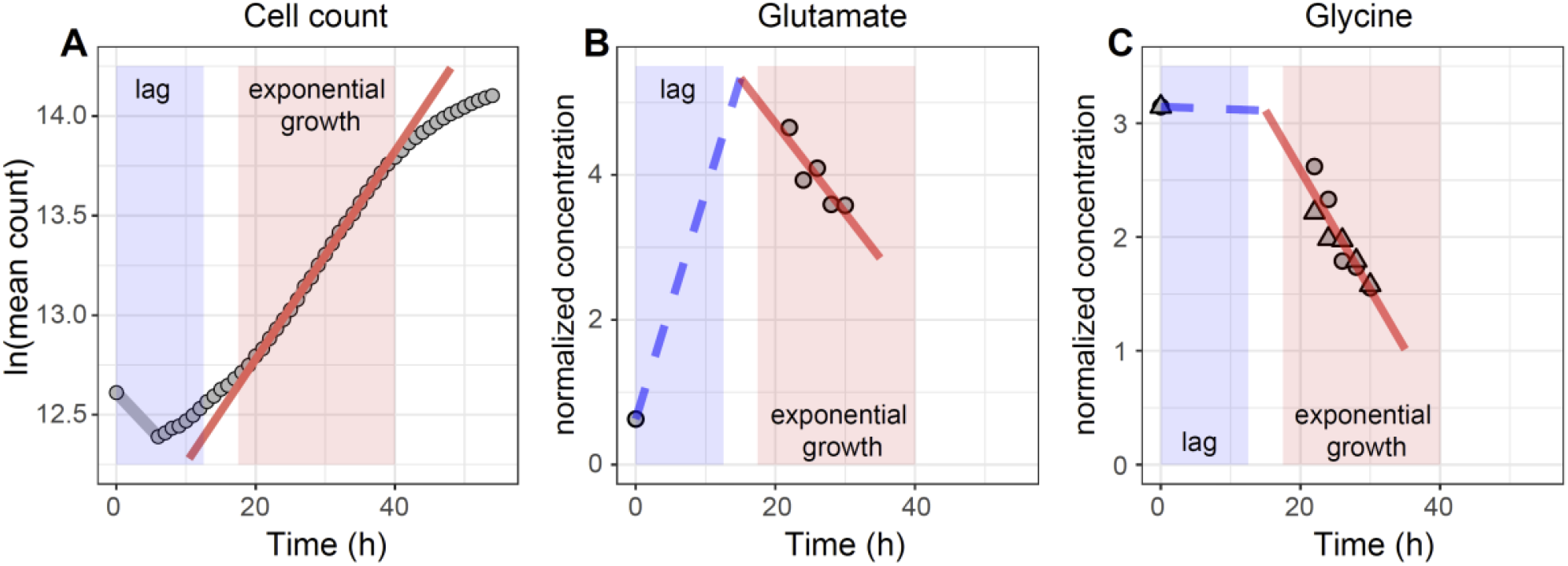
Exo-metabolite exchange fluxes in different growth phases. (A), growth curve of MCF10A cells showing a distinct lag phase (blue box) and an exponential growth phase (red box and red line). (B-C), *r*_*Glu*_ and *r*_*Gly*_ showing distinct metabolic profiles during the lag phase and the exponential growth phase. The REGP method is used to calculate the exchange fluxes during the exponential phase (solid red line). Exchange fluxes in the lag phase is estimated by connecting a straight line from the fresh media sample at t=0 h, to the projected exo-metabolite concentration at t=15 h based on REGP calculations (blue dashed line).

For other nutrients, such as glucose, glutamine, and other amino acids, in general we observed that *r*_*m*,*REGP*_ for nutrient consumption were larger (that is, more negative) than *r*_*m*,*REGP*_, consistent with a higher nutrient consumption rate during exponential growth compared to the lag phase (Fig 1C). For the metabolic waste lactate, we observed that the exchange flux for lactate release was smaller (that is, less positive) by the REGP method (Fig 1C), suggesting that lactate production is elevated during the lag phase, and reduced during exponential growth.

### Simulating steady-state cell growth

A common way to benchmark FBA- and FVA-based GEM simulations is to estimate the cell growth rate (3,15), which can be easily validated experimentally. To do this, we first constructed cell line-specific GEMs by tINIT (16) using cell-line specific transcriptomics data, generated in-house for MCF10A cells (Supplemental Table 2; GEO accession GSE293588) and mined from the Cancer Cell Line Encyclopedia (17) for the 11 cell lines of the NCI-60 panel. We then specified the metabolites that are present in the cell culture media (Ham’s medium), followed by constraining the exo-metabolite exchange fluxes (Fig 1C). For MCF10As, either *r*_*m*,*CORE*_ or *r*_*m*,*REGP*_ were used; for the 11 cell lines of the NCI-60 panel, only the available *r*_*m*,*CORE*_ were used (see Fig 3). Based on these input data as model constraints, the range of feasible *in silico* growth rates were simulated by maximizing and minimizing biomass production. We found that the *r*_*m*,*CORE*_-constrained MCF10A model was infeasible (Fig 3A), meaning that the *in silico* cell was unable to ‘grow’ with the CORE-calculated metabolite uptake and secretion rates. In contrast, the *r*_*m*,*REGP*_-constrained MCF10A model was feasible (Fig 3A). Critically, the experimentally measured growth rate fell within the GEM-simulated solution space (Fig 3A), indicating that GEM-simulations are physiologically relevant when using *r*_*m*,*REGP*_ as constraints, but not with *r*_*m*,*CORE*_. Similar to the *r*_*m*,*CORE*_-constrained MCF10A model, most of the *r*_*m*,*CORE*_-constrained models of the NCI-60 panel cell lines were also either infeasible, or do not contain the experimentally measured growth rate within the simulated solution space (Fig 3A).

**Figure 3.**
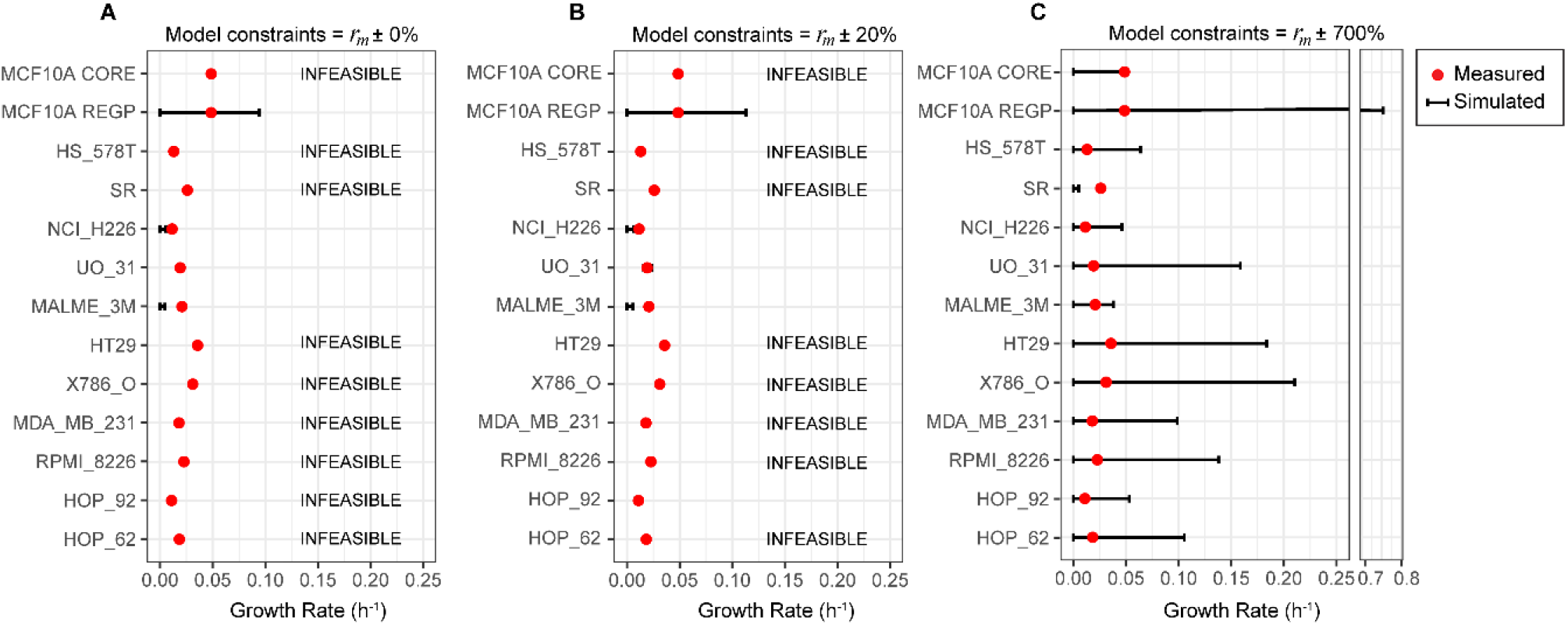
FVA simulation of cell growth rates. (A-C), each cell line-specific GEM model was constrained with the corresponding cell line-specific exo-metabolite exchange fluxes (*r*_*m*_) with a flexibilization factor of 0% (A), 20% (B), and 700% (C). Black bars, FVA-simulated minimum and maximum *in silico* cell growth rate for each cell line. Red dots, experimentally measured growth rate for each cell line.

As a sensitivity analysis, we included a flexibilization factor ranging from 0-20% for all *r*_*m*_ used as model constraints, and found that that this did not play a role in determining model feasibility or the physiological relevance of the simulations (Fig 3A-B). The *r*_*m*,*CORE*_-constrained MCF10A model only produced physiologically relevant simulations when the flexibilization factor was increased to 700%; even at this point, the *r*_*m*,*CORE*_-constrained model of the SR cell line still performed poorly (Fig 3C).

### Organization of the feasible flux space

We then took the *r*_*m*,*REGP*_-constrained MCF10A model as described above, and added a constraint of the biomass production reaction with the experimentally measured growth rate, to produce a constrained GEM of the MCF10A cell line that makes use of all available data. We used this model to explore the feasible flux space of the entire metabolic network of the cell. Following FVA for every metabolic reaction, we calculated the fractional representation of different metabolic subsystems in a sliding window of 200 reactions, ordered by increasing flux variability (Fig 4; Supplemental Table 3). This analysis showed that metabolic subsystems related to fatty acid metabolism, including for example fatty acid biosynthesis pathways and beta oxidation pathways, exhibited low variability in reaction flux (Fig 4). In contrast, amino acid metabolism (AAM) and most central carbon metabolism (CCM) pathways showed intermediate to high levels of variability (Fig 4), even though the exo-metabolite exchange fluxes used as model constraints were all related to CCM (glucose, lactate) and AAM, consistent with previous observations (13,18). For nucleotide metabolism, we observed two distinct regions in this analysis, one with intermediate variability and another with high variability. Finally, we found that reactions in sphingolipid and steroid metabolism, as well as miscellaneous reactions such as pool reactions and artificial reactions necessary for model simulations, exhibited high flux variability (Fig 4).

**Figure 4.**
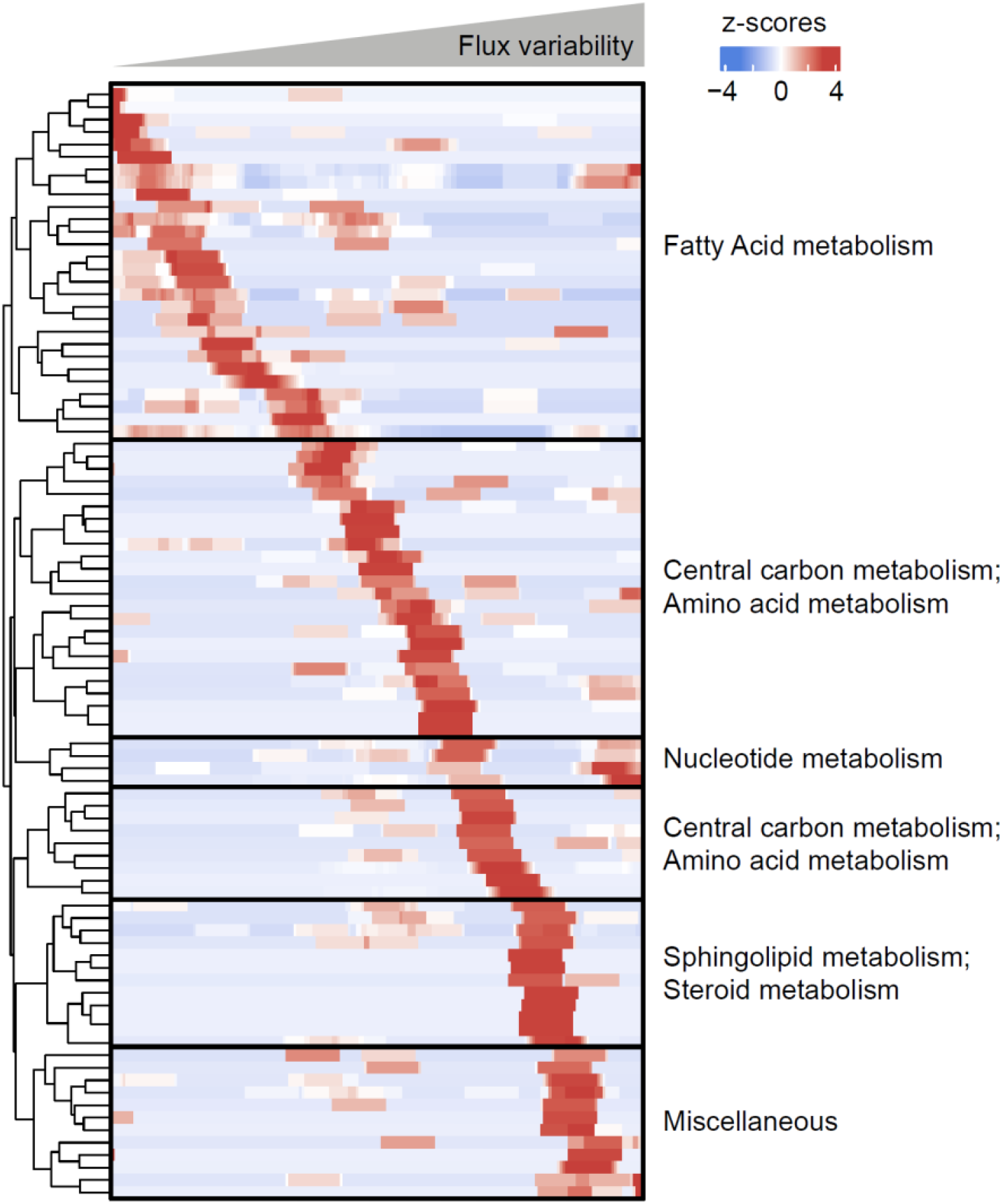
Organization of the feasible flux space of MCF10A cells. The MCF10A-specific GEM was constrained with *r*_*m*,*REGP*_ and the measured growth rate, both with a flexibilization factor of 20%. FVA was performed to determine the flux variability (i.e. the feasible solution space) for every metabolic reaction. The fraction of each metabolic subsystem was calculated in a sliding window of 200 reactions of increasing flux variability, followed by a z-score transformation to facilitate comparison. For ease of visualization, z-scores of > 3 or < -3 were set to 3 or -3, respectively.

## Discussion

Understanding human cell metabolism at the systems level is of critical interest in many areas of health and medicine. GEM-based simulations have shown very promising applications in microorganisms (9–11), but obtaining the necessary input data in human cells has proven to be difficult. A number of methodologies have been developed to leverage transcriptomics data as model constraints (19–23), with mixed results (24), likely because metabolic fluxes are poorly reflected by the abundance of (transcripts of) enzymes in a cell. More recently, exo-metabolite exchange fluxes have been used (3,9,13,25), based on a comparison of exo-metabolite concentrations between the ‘spent’ and ‘fresh’ media (4,13). The caveat of this method is that it violates the steady-state assumption of FBA and FVA, and thus should be used with caution. To address this limitation, here we determined exo-metabolite exchange fluxes by collecting multiple samples exclusively during exponential growth phase (Fig 1B-C). Our results indicated a substantial difference in exchange fluxes in different phases of cell growth (Fig 2), underscoring the importance of distinguishing between growth phases when studying cell metabolism. With the exponential growth-phase exchange fluxes as model constraints, GEM simulations were biologically meaningful, with the measured cell growth rate falling within the simulated solution space (Fig 3). This allowed us to explore the entire metabolic network of the cell with a physiologically relevant flux profile, revealing a distinct organization of the feasible flux space by metabolic subsystems (Fig 4).

Previously, cell-specific GEMs constrained by the exchange fluxes of glucose, lactate, and threonine (all calculated by the CORE method), were shown to predict the cell growth rate to a reasonable degree of agreement with experimentally measured cell growth rates (3). However, incorporation of additional (CORE-calculated) exo-metabolite exchange fluxes, with a flexibilization factor of up to 20%, lead to a large number of infeasible models (Fig 2A-B); suggesting underlying problems with the model constraints and the biological relevance of the simulations. With the REGP method, model simulations remained feasible with a larger number of measured exo-metabolite exchange fluxes, and the feasible flux space was constrained to a biologically meaningful region (Fig 2). We found that the maximum *in silico* growth rate exceeded the experimentally determined growth rate by approximately 2-fold (Fig 3A-B). This suggests that a portion of the consumed nutrients is diverted into non-growth related metabolic tasks, consistent with the notion that, perhaps with the exception of fast-growth cancer cells, human cells do not operate to solely maximize growth (24,26).

Our results show that, even though the input constraints of our model were all related to central carbon metabolism (glucose, lactate) or amino acid metabolism (see Fig 1C), there is nevertheless an intermediate level of flux variability in these subsystems (Fig 4). This is in line with previous work showing that these subsystems do not operate at full capacity in growing cells (18,27). In contrast, we found that reactions in fatty acid metabolism exhibited low flux variability, while reactions in sphingolipid metabolism and steroid metabolism exhibited high flux variability, likely reflecting the degrees of connectivity (i.e. pathway branching) in these subsystems (3,28).

Our findings demonstrate that constraining GEMs with exo-metabolite exchange fluxes calculated by the REGP method allows for accurate model simulations. While this approach demands more resources for exo-metabolite measurements, we believe that it is crucial to profile the metabolic fluxes of human cells at the genome-scale, which can lead to a better understanding of the metabolic process in healthy cells and the identification of potential metabolic targets in diseases.

## Methods

### Cell culture

MCF10A cells were purchased from ATCC (Cat# CRL-10317). Cells were cultured in DMEM/F12 (Cat# 11320033, Gibco) supplemented with MEGM Mammary Epithelial Cell Growth Medium SingleQuots Kit (Cat# CC-4136, Lonza) without GA-1000, along with 0.1 µg/mL cholera toxin (Cat# BML-G117, Enzo Life Sciences) and 100 U/mL penicillin-streptomycin (Cat# 15140, Gibco). Cells were tested for mycoplasma contamination routinely.

### Cell proliferation assays

Absolute cell counts were measured at 22, 26, 30, 46, 50, and 54 h after seeding using the CyQUANT Cell Proliferation Assay kit (Cat# C7026, ThermoFisher Scientific) with a CLARIOstar Plus plate reader (BMG LABTECH). Cell proliferation at all other time points were measured by Incucyte ZOOM (Essen Bioscience), then converted to absolute cell counts based on the corresponding cell counts from the CyQUANT measurements.

### Biomass determination

Cells were harvested with 0.05% trypsin-EDTA (Cat# 25300054, Gibco) and counted using 0.4% trypan blue (Cat# 15250061, Gibco) in a TC20 Automated Cell Counter (BioRad). The cell suspension was transferred into pre-weighed Eppendorf tubes and pelleted by centrifugation at 200 g for 5 mins. Pellets were dried in a microwave at 360 W for 20 mins, and desiccated in a desiccator for >3 days.

### Exo-metabolite measurements

Sampling for exo-metabolites was done during cellular exponential growth phase between 22-30 h after seeding by collection of culture supernatant. Glucose and lactate concentrations were quantified as described before (29), using an HPLC (Shimadzu) with an Aminex HPX-87H column (Cat# 1250140, BioRad) at 65 °C and an IR detector. The column was eluted with 5 mM H_2_SO_4_ at a flow rate of 0.6 mL/min for 26 min. Amino acids were quantified as described before (30), with the aTRAQ Kit (Cat# 4442673, AB Sciex) using a Nexera UHPLC system (Shimadzu) coupled to a Qtrap 6500+ system (AB Sciex) with a BEH C18 column (150 x 2.1 mm, 1.7 μm) (Cat# 186002353, Waters) at 50 °C. A gradient elution of water (eluent A) and methanol (eluent B), both containing 0.1% formic acid and 0.01 % heptafluorobutyric acid, were used as the mobile phases with a constant flow of 1 mL/min. The following MS parameters were used: Curtain Gas: 50; Collision Gas: Medium; IonSpray Voltage: 5500 V; Temperature: 500 °C; Ion Source Gas 1: 60; Ion Source Gas 2: 50. The gradient method was: 2% B from 0–2.5 min, 2% to 40% B from 2.5–3.9 min, held at 40% B until 4.2 min, 40% to 90% B from 4.2– 6.0 min, held at 90% B until 6.1 min, 90% to 2% B from 6.1–8.0 min. Data acquisition and processing were performed using Analyst and MultiQuant 3.0.3 software. Following data QC, exo-metabolite exchange fluxes (*r*_*m*_) were calculated by CORE (4) or REGP (see below).

### Exo-metabolite exchange flux (*r*_*m*_) calculation by REGP

Exo-metabolite exchange fluxes (*r*_*m*_) were determined by calculating the ratio between spent media and unused media samples, then normalizing to the known metabolite concentrations of the cell culture media (DMEM/F12). Exo-metabolite concentrations were normalized for culture volume and cell dry weight, and a linear model was fitted to regress the concentrations against time. *r*_*m*,*REGP*_ is then taken as the slope of the linear model, i.e. the derivative of the fitted metabolite concentration 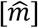 with respect to time *t* (Fig 1B). Goodness of fit was determined by R^2^, with an arbitrary cutoff of 0.7.

### RNA sequencing

RNA was sampled at 27 h after seeding. RNA was extracted using the Quick-RNA Microprep Kit (Cat# R1051, Zymo Research) according to the manufacturer’s protocol. Libraries were prepared from 300 ng RNA with the KAPA RNA HyperPrep Kit with RiboErase (HMR) (Cat# KK8502, Kapa Biosystems). RNA fragmentation was performed for a desired library insert size of 200-300 bp by fragmentation for 6 min at 94 °C. Library concentrations were determined using the Qubit HS kit (Cat# Q32854, Invitrogen). Library size distribution was determined using a High Sensititivy DNA analysis (Cat# 5067-4626, Agilent) on a Bioanalyzer 2100 (Agilent). Libraries with an average between 300-400 bp were loaded on the NextSeq 2000 system (Illumina). Reads were quality controlled, mapped to the human genome hg38, and counted by Seq2science (31), available at https://github.com/vanheeringen-lab/seq2science.

### Genome-scale metabolic modeling

The consensus Human-GEM, Human1 v1.12.0 (3), was used for all procedures detailed below. Human-GEM is a ‘generic’ model which contains all observed metabolites and reactions in human cells. For each cell line, contextualized models were constructed using tINIT (16), where the generic Human-GEM is pruned in a cell line-specific manner based on whether or not the (transcript of) an enzyme is expressed. We used transcriptomics data measured in-house (MCF10As) or mined from CCLE (17), with an arbitrary expression level cutoff of 1 TPM. Each cell line-specific model was first constrained with components of the growth media (without specifying the exchange fluxes), and then constrained by the measured exchange fluxes with a flexibilization factor ranging from 0% to 700%. For simulation of growth rate, the range of feasible *in silico* growth rate was simulated by sequentially minimizing and maximizing the biomass reaction, as implemented in the COBRA toolbox (32). The MCF10A-specific model was further constrained with the measured growth rate to perform FVA for all reactions. The flux variability for each reaction, i.e. max(flux) – min(flux), is sorted from lowest to highest. A sliding window of 200 reactions with a step size of 10 reactions was used to assess the variation in reaction flux across different subsystems. Z-scores were calculated for the fraction of reactions per subsystem in a given window. Exchange reactions, transport reactions, and subsystems with less than 5 reactions, were excluded from the visualization of this analysis in Fig 4. All simulations were performed using MATLAB 2023a (MathWorks, Inc.) with Gurobi solver v10.0.1 (Gurobi Optimizer).

## Data and code availability

Raw RNAseq data are available at GEO, accession GSE293588. All other data and code used in this paper are available in the GitHub repository (https://github.com/Radboud-YuLab/FluxProfilingREGP). Procedures related to metabolic modeling are implemented in MATLAB (2023a). Numerical analyses and graphics are done in R v4.2.3.

## Acknowledgements

The Yu lab is supported by grants from the Dutch Cancer Society and the Radboud-Western Collaboration Fund. We thank Niky Thijssen for technical support during the experimental work.

## Author contributions

Conceptualization: RY. Data curation: CH, XJ. Formal analysis: CH, XJ. Funding acquisition: YC, RY. Supervision: YC, RY. Writing – original draft: CH. Writing – review and editing: XJ, YC, RY.

